# The common kestrel *Falco tinnunculus* is attracted to the scent of voles

**DOI:** 10.64898/2025.12.02.690931

**Authors:** Luisa Amo, Concepción Salaberria, Ana Martín Silva, Ignacio Otero Cañas, Juan Antonio Fargallo

## Abstract

Although the study of the role of olfaction in avian foraging is providing exciting results, whether olfaction is involved in the detection of prey by raptors is still understudied. Here, we present the results of a study aimed to assess whether the common kestrel (*Falco tinnunculus*) is attracted to the scent of the cues that common voles (*Microtus arvalis*) use to mark their territory. In an experimental study in captivity we released kestrels in an aviary containing two trays, one tray containing vole scent and one unscented tray. Results show that kestrels are attracted to the vole chemical cues as a greater number of kestrels paid the first visit to the vole-scented tray. Furthermore, the kestrels visited significantly more often the vole scented tray than the control tray. Therefore, our results show that common kestrels are able to use chemical cues emitted by their prey for foraging.

## Introduction

The most ancient mechanism used by organisms to assess their environment, including searching for prey, is chemoreception [1]. The use of chemical cues in foraging has been extensively studied in different taxa but to a lesser extent in birds, despite foraging strategies have played a fundamental role in the evolution of chemosensory abilities in birds [2]. Empirical studies have shown the importance of olfaction in finding prey in predatory birds [3,4]. At sea, Procellariformes and Sphenisciformes seabirds are able to recognise dimethyl sulphide, a chemical compound signalling areas of high productivity where they can encounter their prey [5,6]. Olfaction also plays an important role in plant–herbivory–predator interactions in terrestrial systems. Birds detect volatile compounds released by trees when attacked by herbivorous arthropods which favours their predation [7], as insectivorous birds exploit the herbivore induced volatiles emitted by plants in response to caterpillar herbivory [3,8]. Insectivorous birds not only use indirect cues to find their prey, but are also able to detect the chemical cues emitted by the prey itself. Insectivorous birds use olfaction to detect the pheromones of adult Lepidoptera and use these to locate their prey [9].

Although the study of the role of olfaction in avian foraging is providing exciting results, whether olfaction is involved in the detection of prey by raptors is still understudied [10]. First studies that showed evidence of the use of olfaction in raptor foraging were provided in scavengers such as the Turkey vulture (*Cathartes aura*, [10, 11, 12]) or the greater yellow-headed vulture (*Cathartes melambrotus*, [13]). Outside of the group of scavengers, only in one species that feeds on pollen, the honey buzzard (*Pernis orientalis*), has food detection been observed through smell [14]. However, it is still unknown whether olfaction is involved in foraging in birds of prey that consume mobile prey (orders Accipitriformes and Falconiformes), as they have been considered to rely principally on vision. Previous evidence suggests that some predatory species of raptors, such as the common kestrel (*Falco tinnunculus*), are visually attracted to the UV light reflected by the urine and faeces marks of their small mammal prey [15, 16]. However, first suggestion that this detection of prey may not depend entirely on UV vision came from the study of Zampiga and collaborators [17], as they found that adult kestrels also prefer scent-marked areas under UV-filtered light. Furthermore, later studies analysing raptor UV sensitivity suggested that vole urine is unlikely to provide a reliable visual signal to hunting raptors [18]. Therefore, other cues present in urine, such as smell, might be important in the detection of prey areas or individuals in birds of prey.

The common kestrel is a small raptor species that has been traditionally considered a vole specialist in northern latitudes [19], while its diet is made up of a wide range of vertebrate and invertebrate prey species in more southern latitudes [20], although common voles (*Microtus arvalis*) remains the preferred prey even in more diverse habitats [21]. Here, we present the results of a study aimed to disentangle whether birds of prey use olfaction to find their prey. Concretely, we performed an experimental study to examine whether the common kestrel is attracted to the chemical cues that common voles use to scent-mark the territory.

## Material and methods

### Study species

The kestrels used in the present study come from a rehabilitation centre in Madrid province (GREFA). The birds were brought to the centre in order to be recovered, as they were captured as fletching birds or adults that had any health problem (e.g., collisions with cars, windows). Before the birds were released back to nature, they were tested in our experiment some days before. Therefore, the birds were healthy and able to search for food by themselves. We used 38 birds, 11 adults and 27 juveniles. The birds were located in indoor aviaries (3 × 5 × 3 m) that had an outdoor part. They were fed once a day with one day old chickens and they had access to water ad libitum.

### Experimental design

The study was performed in August and September in 2020 and 2021 in two indoor aviaries (3 × 3 × 3 m) in GREFA installations. The aviaries had 2 wooden perches. There were two trays on the floor, one under each perch, containing papers with the corresponding treatment. Trays (54 × 40 × 9 cm) were made of polypropylene. The experimental tray contained absorbent papers soiled with prey scent of one vole, and the other tray, the control tray, contained absorbent papers with some drops of water, to have an odourless treatment with similar level of humidity than the experimental treatment. The trays were covered with shading fabric to avoid kestrels visually detecting the treatments. The location of the treatments was randomized between trials. The prey scent treatment was located 17 times under the left perch and 21 times under the right perch. The trays were separated 2 meters.

The vole scent was obtained from three male common voles captured in Campo Azálvaro (Segovia, central Spain; [21]). The voles were captured with Sherman traps baited with a mixture of tuna, flour and oil and with a piece of apple and transport to the lab in such traps. The voles were housed in individual cages (50 × 40 × 40 cm) during one moth to obtain absorbent papers soiled with prey scent. They were fed daily with vegetables and had water ad libitum. After that, they were released in the exact place of capture. We obtained vole scent by placing absorbent papers (49 × 39 cm filter paper made of cellulose fibres) in the individual cages of voles during 3 days. After that, the papers were replaced and the soiled absorbent papers were stored in polyethylene plastic bags and frozen at – 20 °C until they were used in the experiments.

The kestrels were fed the previous day, so they were at least 20 hours without food when they were tested. Trials were performed between 9:00 and 14:00 hours. One experimenter captured the focal kestrel in the aviary and brought it inside a transport cage to the experimental aviary, where it was released. The behaviour of the kestrel was recorded for 2 hours with a video camera that was located in one of the walls of the aviary, close to the roof. After two hours, an experimenter captured the kestrel and returned it to its aviary. Birds were tested only once. After each trial, the trays, the perches and the floor of the experimental aviary was carefully cleaned with 70° alcohol.

After the trials, an experimenter, blind to treatments, analysed the kestrel behaviour and recorded the first tray visited and the number of visits to each tray.

### Statistical analysis

We defined two variables to analyse kestrel behaviour: 1) “first tray visited”, describing which tray (the smelly or the control) was approached first, and 2) “proportion of visits to the prey scented tray”, which is the number of visits to the experimental tray divided by the total number of visits to both trays. The first tray visited and the proportion of visits to the experimental tray were analysed by using generalised linear models fit by the Laplace approximation, with the dependent variables following a binomial distribution with logit function. The kestrel age (juvenile vs adult) and the tray location (left vs right) were included in the initial models as fixed factors and the aviary as a random factor. Non-significant factors were removed from the final models. Analyses were performed with the Statistical package R 4.4.2 [22].

## Results

Ten kestrels (2 adults and 8 juveniles) did not visit any tray and were excluded from the analysis. Therefore, the statistical analysis was performed with 28 kestrels (10 adults and 18 juveniles).

The location of the tray influenced first tray visit (*Z* = - 2.48, *df* = 1, *p* = 0.01, 95% CI [0.003, 0.35]). More birds paid the first visit to the vole scented tray when it was located on the left. Controlling for this influence, the model showed differences in the number of kestrels that paid the first visit to the vole scented tray or the unscented tray (*Z* = 2.09, *df* = 1, *p* = 0.04, 95% CI [0.55, 0.99]), with more birds paying the first to the vole scented tray than to the control unscented tray. The age of the kestrels did not influence the first visit and was removed from the final model (Table 1).

**Table 1.** Initial model of the analysis of first tray choice, i.e. the difference in the number of kestrels (n = 28) that visited the tray in relation to the scent treatment (prey scented tray vs control odourless tray). The model included the tray location (right vs left) and the age of kestrels (adult vs juvenile) as fixed factors.

|  | Estimate | Std. Error | Z | P |
| --- | --- | --- | --- | --- |
| Treatment | 2.45 | 1.11 | 2.22 | 0.03 |
| Location | -2.57 | 1.20 | -2.14 | 0.03 |
| Age | -1.54 | 1.05 | -1.47 | 0.14 |

The location of the tray also influenced the proportion of visits, with birds visiting the tray located on the left significantly more often than the tray on the right (*Z* = -2.53, *df* = 1, *p* = 0.01, 95% CI [0.21, 0.46]). Controlling for this influence, the model showed that the kestrels visited the vole-scented tray significantly more often than the control tray (*Z* = 2.77, *df* = 1, *p* = 0.006, 95% CI [0.55, 0.77], Fig. 1). The age of the kestrels did not influence the proportion of visits to the trays and was removed from the final model (Table 2).

**Table 2.** Initial model of the analysis of the percentage of visits of common kestrels (n = 28) to the tray in relation to the treatment (prey scented tray vs control odourless tray). The model included the tray location (right vs left) and the age of kestrels as fixed factors.

|  | Estimate | Std. Error | Z | p |
| --- | --- | --- | --- | --- |
| Treatment | 0.65 | 0.26 | 2.46 | 0.01 |
| Location | -0.75 | 0.29 | -2.55 | 0.01 |
| Age | 0.13 | 0.27 | 0.48 | 0.63 |

**Fig. 1.**
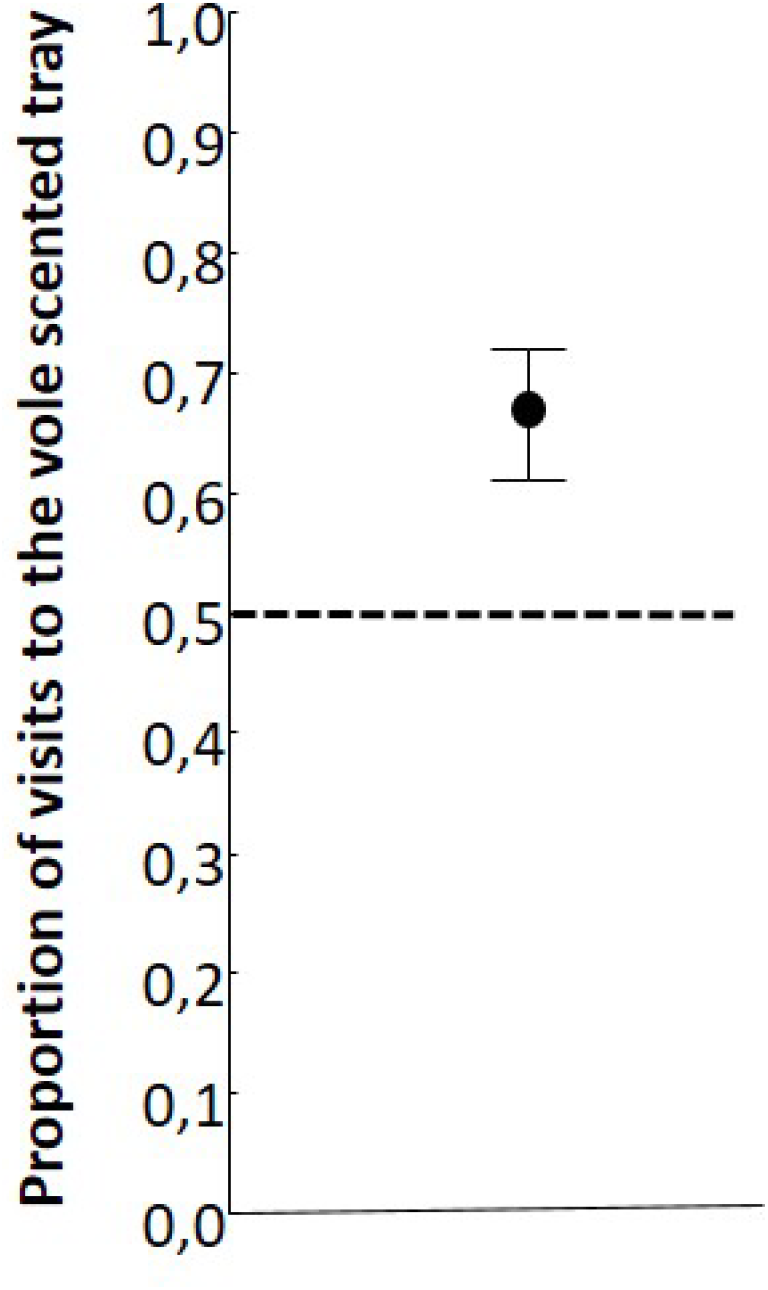
Estimated proportion of visits to the experimental tray that contained common vole (*Microtus arvalis*) scent by common kestrels (*Falco tinnunculus*) (*n* = 28), when released in an aviary with two trays under two perches: one control tray (unscented) and one experimental tray (vole scent marks), controlling for the location of the tray. Trays were covered with shading fabric that allowed detection of chemical cues but avoid visual detection.

## Discussion

The kestrels detected the chemical cues of their main prey, the common vole, and they were attracted to vole scent marks. More birds paid the first visit to the vole-scented tray than to the control tray. Furthermore, the birds visited the tray with scent-marks significantly more often than the tray with the odourless control. Our experimental study guarantees that kestrels were only attracted to chemical cues and not visual cues, because papers containing the vole chemical cues were covered by an opaque shading fabric, preventing birds to see throughout it. Therefore, we can exclude that birds were attracted to the UV cues emitted by vole scent marks. Previous evidence suggested that common kestrels could be visually attracted to the UV light reflected by the urine and faeces marks of their small mammal prey [15, 17, 23]. However, our results agree with previous results from other studies indicating that this detection of prey may not depend entirely on UV vision, because adult kestrels were also attracted to scent marks even with a UV filter [17] and because the UV light emitted by vole urine is outside the UV light detection range of kestrel eye [18].

We randomized the side of the aviary where the tray containing vole scent was located. However, despite this randomization, we found that the location of the tray influenced the first choice as well as the percentage of visits of the scented tray, with more birds visiting for the first time the vole-scented tray when it was on the left side. The birds also visited more frequently the experimental tray when it was located on the left side. The influence of the location of the tray in the behaviour of birds may be explained by the fact that, despite the location of the treatment was initially randomized between trials, after removing the birds that did not visit any tray, the location of the prey scent was more often on the left (18) than on the right (8). Furthermore, previous evidence suggests that common kestrels exhibit brain lateralization, displayed in the direction of body rotation when perceiving a sound stimulus from behind the body [24]. Kestrels exhibited an anticlock movement of the body, i.e. a left preference [24]. Our results suggest that this brain lateralization seems also to be exhibited in response to prey scents. Anyhow, because we controlled for the location effect in the analysis of percentage of visits to both trays, and found that the kestrels visited the vole-scented trays significantly more often, we are confident that the kestrels were attracted to prey chemical cues.

Adult and juvenile kestrels were similarly attracted to the scent of vole chemical cues. This result is in accordance with previous results of Zampiga and collaborators [17] that demonstrated that the attraction to vole scent marks has an innate component in kestrels, as naïve juveniles were attracted to vole scent marks under natural light but not with a UV filter [17]. The innate attraction to vole scent suggests that this ability is under a strong selection pressure, as juvenile kestrels are not taught hunting by their parents [25] and they may be able to hunt without parental help between 2-4 weeks after fledging [19]. Innate detection of prey chemical cues has also been shown in avian species where parents do not teach prey capture to offspring, such as Procellariformes and Sphenisciformes, where naïve nestlings are attracted to DMS [6, 26]. In contrast, in insectivorous birds, where parental care islonger and parents taught fletching searching for food, attraction to prey-associated scents (HIPVs) is not innate and may be learned [27, 28].

Research has previously demonstrated that the rough-legged buzzard (*Buteo lagopus*; [23]) and the great grey shrike (*Lanius excubitor*; [29]) are attracted to prey scent marks during hunting. As with kestrels, this behavior was thought to be mediated by the visual detection of the marks. Our results with kestrels, however, raise the possibility that buzzards and shrikes can also detect the volatiles released from these scent marks.

In conclusion, our study shows that raptor species that hunt for mobile prey are sensitive to the chemical signals emitted by the prey. The short-range detection observed in our study gives the chance of raptors to detect areas with the presence of prey species, which in colonial species, such as many vole species, with high densities of individuals, can be detected at long distances, a beneficial foraging strategy similar to that used by other bird species sensitive to dimethyl sulphide or HIPVs that indirectly reveal the presence of prey. Our results also add to the evidence that the scent marking of the territory may increase predation risk to small mammals not only by mammalian predators but also by diurnal raptors.

## Acknowledgment

We especially thank Jorge Aguado and GREFA personnel for the use of their facilities.

## Ethicals

The experiment was carried out under license of the Consejería de Medio Ambiente, Agricultura e Interior of Comunidad Autónoma de Madrid (PROEX 261/19).

## Author contributions

C. S. collected data, A.M.S. and I.O.C. help with data collection, L.A. wrote the first draft of this manuscript, C.S., L.A. and J.A.F. were responsible for the experimental design, statistical analysis and contributed significantly in later drafts of the paper.

